# Single-molecule imaging of mRNA localization and regulation during the integrated stress response

**DOI:** 10.1101/332502

**Authors:** Johannes H. Wilbertz, Franka Voigt, Ivana Horvathova, Gregory Roth, Yinxiu Zhan, Jeffrey A. Chao

**Author notes:** Dr. Jeffrey A. Chao, office: Maulbeerstrasse 66, CH – 4058 Basel.

## Abstract

Biological phase transitions form membrane-less organelles that generate distinct cellular environments. How molecules are partitioned between these compartments and the surrounding cellular space and the functional consequence of this localization is not well understood. Here, we report the localization of mRNA to stress granules(SGs) and processing bodies(PBs), which are distinct biomolecular condensates, and its effect on translation and mRNA degradation during the integrated stress response. Using single mRNA imaging in living human cells, we find that the interactions of mRNAs with SGs and PBs have different dynamics and that specific RNA binding proteins can anchor mRNAs within these compartments. During recovery from stress, mRNAs that were within SGs and PBs are translated and degraded at similar rates as their cytosolic counterparts.

---

Activation of the integrated stress response results in a global inhibition of translation that coincides with the appearance of cytosolic membrane-less organelles known as stress granules (SGs) and processing bodies (PBs) formed by the condensation of RNAs and RNA-binding proteins. While considerable effort has been invested in characterizing the biophysical properties that govern the formation of these granules, the molecular mechanisms attributed to their regulatory function are not completely understood (*1*). The prevailing models for the function of granules have been built upon observations of the behavior of their protein constituents and, consequently, the effects on the bound RNAs have often only been inferred (*2, 3*). Recently, methods have been devised for the isolation of SGs or PBs from mammalian cells, which has allowed the RNA content of these granules to be identified, however, these approaches are unable to investigate the underlying dynamics of these interactions (*4–8*). Here, we used single mRNA imaging in living cells to directly monitor the spatial and temporal localization and regulation of mRNAs during the integrated stress response.

We engineered a HeLa cell line expressing mRNAs, PBs, and SGs labeled by three spectrally distinct fluorophores to allow their simultaneous detection in living cells (Fig. 1A). First, G3BP1-GFP and DDX6-TagRFP-T were stably integrated into HeLa cells and served as SG and PB markers, respectively. Cells were then sorted for low GFP and TagRFP-T levels by fluorescence activated cell sorting(FACS) to prevent SG or excess PB formation in the absence of stress (*9*). After generation of this cell line, the cells were treated with 100 *μ*M sodium arsenite (SA) to confirm that eIF2α was phosphorylated on Ser51 and that translation was inhibited indicating the activation of the integrated stress response (fig. S1A,B). The number of G3BP1-GFP and DDX6-TagRFP-T granules was similar to the levels observed for endogenous G3BP1 and DDX6 in the presence of SA (fig. S1C). Next, we confirmed that the size, number, and formation dynamics of both G3BP1-GFP and DDX6-TagRFP-T granules were comparable with previous reports (fig. S1D,E) (*10, 11*).

**Fig. 1:**
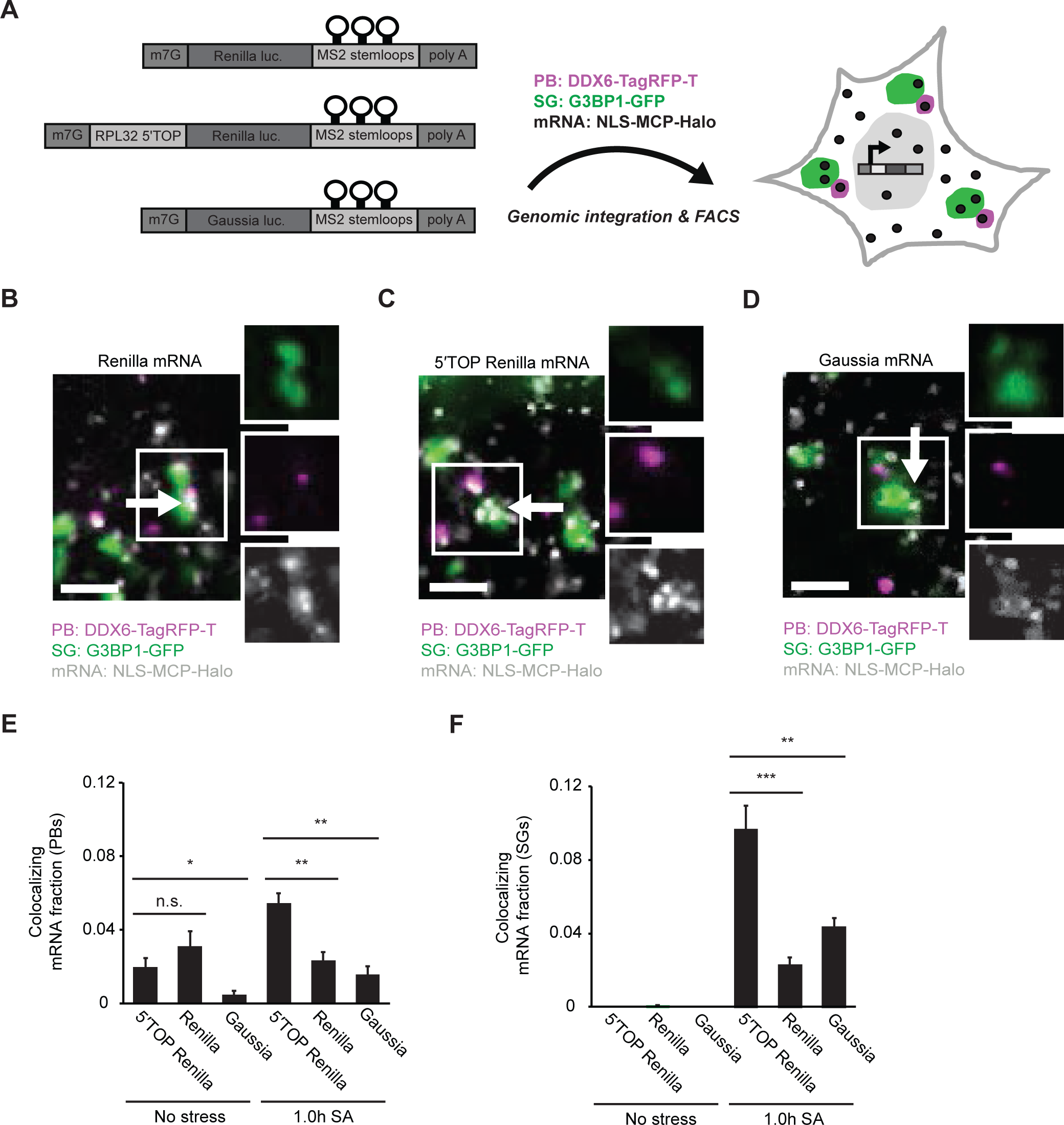
Three-color live cell imaging identifies distinct mRNA localization to PBs and SGs. (A) Schematic depicting mRNA reporters and HeLa cell line expressing DDX6-TagRFP-T (PBs) and G3BP1-GFP (SGs). mRNAs were expressed from a doxycycline inducible single locus and labelled with NLC-MCP-Halo. (B) Representative image of localization of Renilla reporter mRNAs (white) to PBs (magenta) and SGs (green) during stress. (C) Representative image of localization of 5′TOP Renilla mRNA reporters (white) to PBs (magenta) and SGs (green) during stress. (D) Representative image of localization of Gaussia reporter mRNAs(white) to PBs(magenta) and SGs(green) during stress. (E,F) Colocalization analysis and quantification of the data presented in (B-D). All mRNA reporters localized to PBs (E) and SGs (F) during SA stress, but 5′TOP Renilla reporter mRNAs were significantly more enriched than Renilla or Gaussia mRNA reporters. Arrows indicate mRNA colocalization with SGs. Scale bars = 2 µm; mean ± SEM; two-tailed, unpaired Student′s t-test; * = p < 0.05, ** = p < 0.01, *** = p < 0.001; > 20 fields of view per time point and experiment, 3 biological replicates.

To detect mRNAs in living cells we cloned 24 MS2 stem-loops into the 3‘-untranslated region (UTR) of three different transcripts that we anticipated could have potentially different localization during the stress response (Fig. 1A). The first reporter mRNA contained Renilla luciferase in the coding sequence and was generated to represent a standard mRNA encoding a cytosolic protein. The second mRNA reporter was identical except for the addition the first 50 nucleotides of the RPL32 5UTR which contains a 5′ - terminal oligopyrimidine (TOP) motif. 5′TOP motif-containing ribosomal proteins and translation factors are highly abundant, and are thought to constitute ∼20% of all transcripts present in cells (*12, 13*). Based on previous observations by us and others, we expected the 5′TOP Renilla reporter to accumulate to a greater extent in SGs and PBs compared to the Renilla reporter (*14, 15*). Since earlier reports suggested that ER localization protects mRNAs from entering SGs, the third reporter contained the secreted Gaussia luciferase in the coding sequence and was generated to represent an mRNA that is translated on the endoplasmic reticulum(ER) (*16-18*). Accurate detection and tracking of single mRNA molecules is facilitated by physiological expression levels. Therefore, we stably integrated single-copies of the reporters into a defined genomic locus in doxycycline-inducible HeLa cells (*19*). To visualize mRNAs, we stably co-expressed nuclear localization signal (NLS) containing Halo-tagged MS2 bacteriophage coat protein (NLS-MCP-Halo) that binds with high affinity to MS2 stem-loops (*20–22*). Together, this allowed us to image single mRNA molecules in live unstressed and stressed human cells.

After doxycycline induction, we imaged all three cell lines in the absence and presence of SA. In the absence of stress, Renilla reporter mRNAs rapidly moved throughout the cytosol, SGs were absent and PB numbers were low (fig. S1C, Movie S1). After 1 hour of SA treatment, the majority of Renilla mRNAs still diffused freely in the cytosol, but a fraction of molecules localized to SGs and PBs, which reduced their mobility (Fig. 1B, Movie S2). In unstressed cells, the 5′TOP Renilla reporter mRNAs behaved similar to the Renilla reporter, however, a larger fraction of 5′TOP Renilla reporter mRNAs was localized to SGs and PBs during stress (Fig. 1C, Movie S3 and S4). In the absence of stress, Gaussia mRNA reporters were mostly static in the cytosol, which is consistent with a previous study that demonstrated their translation-dependent association with the ER (*18*) (Movie S5). Upon addition of SA, when translation initiation is inhibited, the majority of Gaussia reporters became mobile. Interestingly, a small fraction of Gaussiam mRNAs did localize to SGs and PBs indicating that ER-association prior to stress does not prevent their entry into granules (*16, 17*) (Fig. 1D and Movie S6)). Single molecule tracking and quantification of mRNA reporter colocalization with PBs and SGs demonstrated that the 5′TOP Renilla reporter mRNAs localized significantly more to both granules than either Renilla or Gaussia mRNA reporters (Fig. 1E,F).

In order to confirm the localization patterns observed in living cells, we performed single molecule fluorescence *in situ* hybridization (smFISH) in HeLa cells with probes against the endogenous GAPDH and RPL32 transcripts, combined with IF against endogenous G3BP1 and DDX6 (fig. S2A,B). Upon addition of SA, only a small fraction of GAPDH transcripts colocalized with PBs and SGs, which is similar to previous reports (*6*) (fig. S2A,C). Endogenous RPL32 transcripts accumulated in PBs and SGs similar to the levels we observed for the 5′TOP Renilla reporter (fig. S2B,C). Taken as whole, our results demonstrate that *cis-*acting elements within transcripts can promote their association with granules during stress.

After having observed the differential localization of Renilla and 5′TOP Renilla mRNA reporters to SGs and PBs, we next sought to understand how this pattern was established. In principle, the differential recruitment of mRNAs to stress-induced mRNPs could either occur during the formation of granules or only after mature granules had formed. To address this question, we quantified the co-localization of Renilla and 5′TOP Renilla transcripts with SGs and PBs over time (Fig. 2A). For PBs we observed that 5′TOP Renilla reporters entered these structures mainly during the first 30 minutes, after which the colocalizing mRNA fraction stayed constant until the end of the time course (Fig. 2B). In contrast, the Renilla reporter showed a significantly smaller time-dependent colocalization increase with PBs (Fig. 2B). mRNA recruitment kinetics to SGs were similar to the results obtained for PBs. 5′TOP Renilla reporters entered SGs faster and in higher numbers than the Renilla transcripts (Fig. 2C). Most mRNAs were recruited during the first 30 minute of SA stress, reaching a plateau phase afterwards. Renilla reporters showed only a modest increase in SG colocalization over time which was significantly smaller than the increase observed for 5′TOP Renilla mRNAs (Fig. 2C). These results demonstrate that 5′TOP-dependent mRNA localization to PBs and SGs correlates with PB and SG formation during stress onset.

**Fig. 2:**
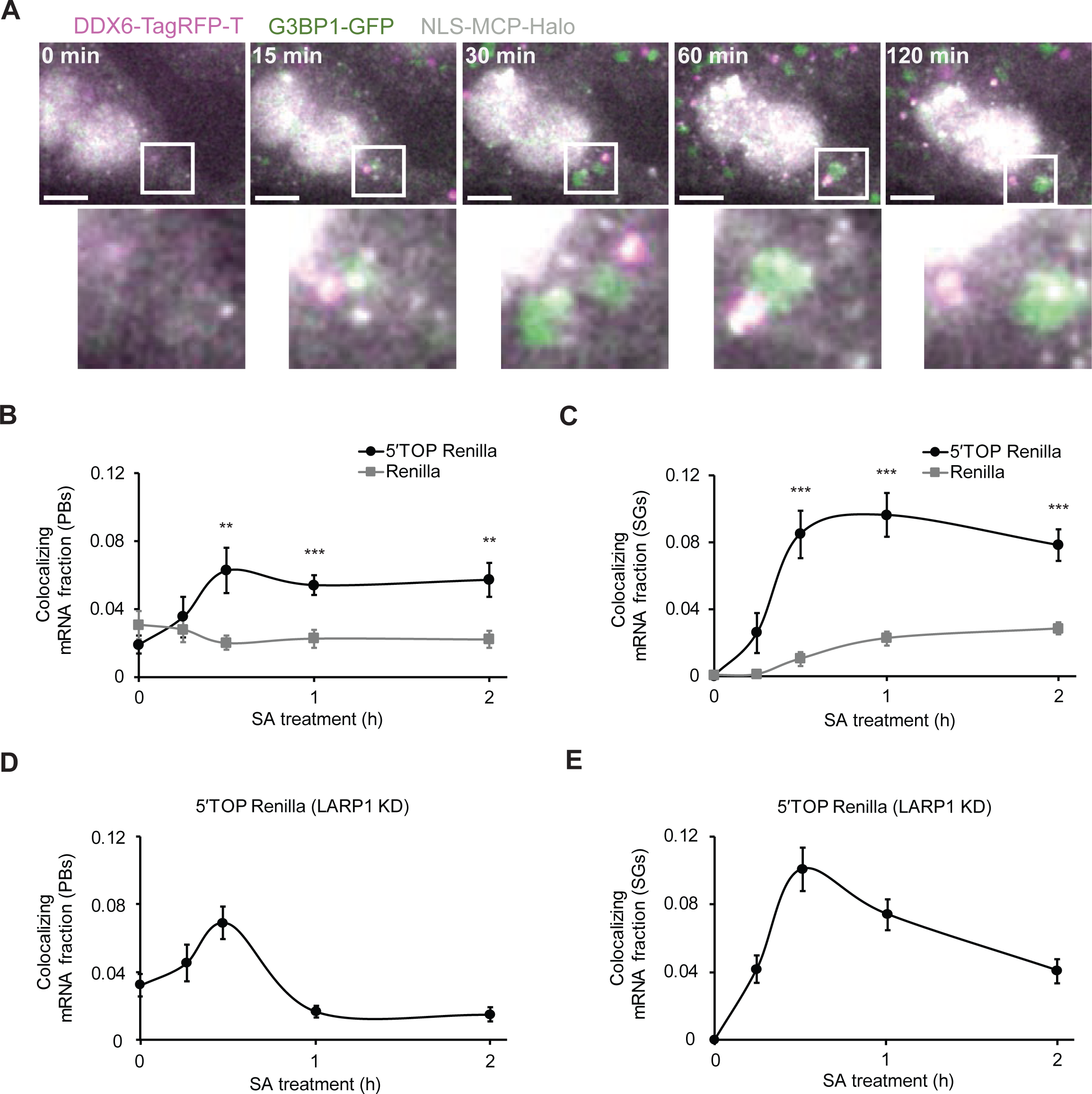
5′TOP mRNA localization correlates with PB and SG formation during stress onset and is LARP1-dependent. (A) HeLa cells stably expressing G3BP1-GFP, DDX6-TagRFP-T, NLS-MCP-Halo and 5′TOP Renilla reporter mRNAs were treated with 100 µM SA for 2 hours and single cells were imaged at the indicated intervals. Cytosolic mRNAs (white) dynamically bound to PBs(magenta) and SGs(green) during and after their formation. (B,C) HeLa cell lines stably expressing G3BP1-GFP, DDX6-TagRFP-T, NLS-MCP-Halo and either Renilla or 5‘TOP Renilla reporter mRNAs were treated with 100 µM of SA for 2 hours and cells were imaged over time and mRNA colocalization with PBs (B) and SGs (C) was assessed. 5′TOP Renilla reporter mRNAs were recruited more to PBs and SGs. (D) and (E) HeLa cell lines stably expressing G3BP1-GFP, DDX6-TagRFP-T, NLS-MCP-Halo coat proteins and 5′TOP Renilla reporter mRNAs were transfected with siRNAs against LARP1 for 48h and treated with 100 µM SA for 2 hours. Cells were imaged over time and the mRNA fraction colocalizing with PBs (D) and SGs (E) was analyzed. Scale bars = 10 µm; mean ± SEM; two-tailed, unpaired Student′s t-test; ** = p < 0.01, *** = p < 0.001; > 20 fields of view per time point and experiment, 3 biological replicates.

Since we observed that the 5′TOP sequence promoted mRNA localization to granules during the stress response, we then asked if there was a *trans-*acting factor that also contributed to this effect. Recently, the RNA binding protein La-related protein 1 (LARP1) has been shown to bind the m^7^G-cap and 5′TOP-element of mRNAs and to regulate their translation (*23–27*). In addition, LARP1 is present in SGs and PBs (fig. S3A) (*28–30*). We decreased levels of LARP1 in HeLa cells by 48h siRNA-mediated knock-down (KD) and performed a 120-minute time-course experiment identical to the one described above (fig. S3A,B). Importantly, LARP1 KD did not affect mRNA numbers as detected by single molecule imaging (fig. S3C). Furthermore, LARP1 KD also did not alter the size or numbers of SGs, while PBs where slightly reduced in size (fig. S3D,E). Interestingly, the association of 5′TOP Renilla mRNAs into PBs and SGs during the first 30 minutes of SA stress was unperturbed (Fig. 2D, E). At later time points, however, the fraction of 5′TOP Renilla mRNA in both granules was reduced demonstrating that LARP1 was necessary for anchoring 5′TOP Renilla within granules. In order to confirm that LARP1 also affected the localization of endogenous transcripts during stress, we performed IF against G3BP1 and DDX6 in combination with smFISH against either RPL32 mRNA or GAPDH mRNA (fig. S4A). RPL32 mRNA localization to SGs, but not PBs, was reduced during LARP1 KD while GAPDH localization to granules was unaffected (fig. S4A,B). These experiments indicate that in addition to granule size and mRNA length, RNA-binding proteins can control the localization of specific transcripts to SGs and PBs during stress and that this regulation can occur even after transcripts have already entered phase separated compartments (Moon et al).

In order to characterize the dynamics of mRNA localization during stress, we extracted directionality information from mRNA tracks relative to PBs and SGs (Fig. 3A). mRNA molecules that were overlapping with a PB or SG received a localization index value of 1 and mRNAs outside of granules received a value of 0. A change of localization index value within one mRNA track therefore indicated a change of direction relative to the granule. This analysis allowed us to distinguish four different categories of mRNA movement relative to PBs and SGs (Fig. 3A). mRNAs could either be classified as static during the observation period, they could show multiple transient interactions, or simply move inside or outside of a granule. Renilla reporters had lower levels for all interactions with PBs and SGs than 5′TOP Renilla reporters (Fig. 3B,C). In addition, no single movement class was significantly more prominent than the others. For the 5′TOP Renilla reporter, it was interesting to see that mRNAs behaved differently when interacting with either PBs or SGs (Fig. 3D,E). Up to half of 5′TOP Renilla reporter localization behavior to SGs was explained by static mRNA interaction with the SG, while the other half was mainly composed of multiple transient interaction and, to smaller extend, unidirectional movements (Fig. 3E). The 5′TOP Renilla reporter interaction patterns with PBs were less dynamic. Here, between 70-85% of the localization behavior was explained by static mRNA interaction with PBs. The remaining fraction was composed of similar amounts of transient and unidirectional movement (Fig. 3D). Note that the distribution of movement patterns was similar at all time points indicating that interactions between granules and RNAs was constant throughout granule formation and maturation.

**Fig. 3:**
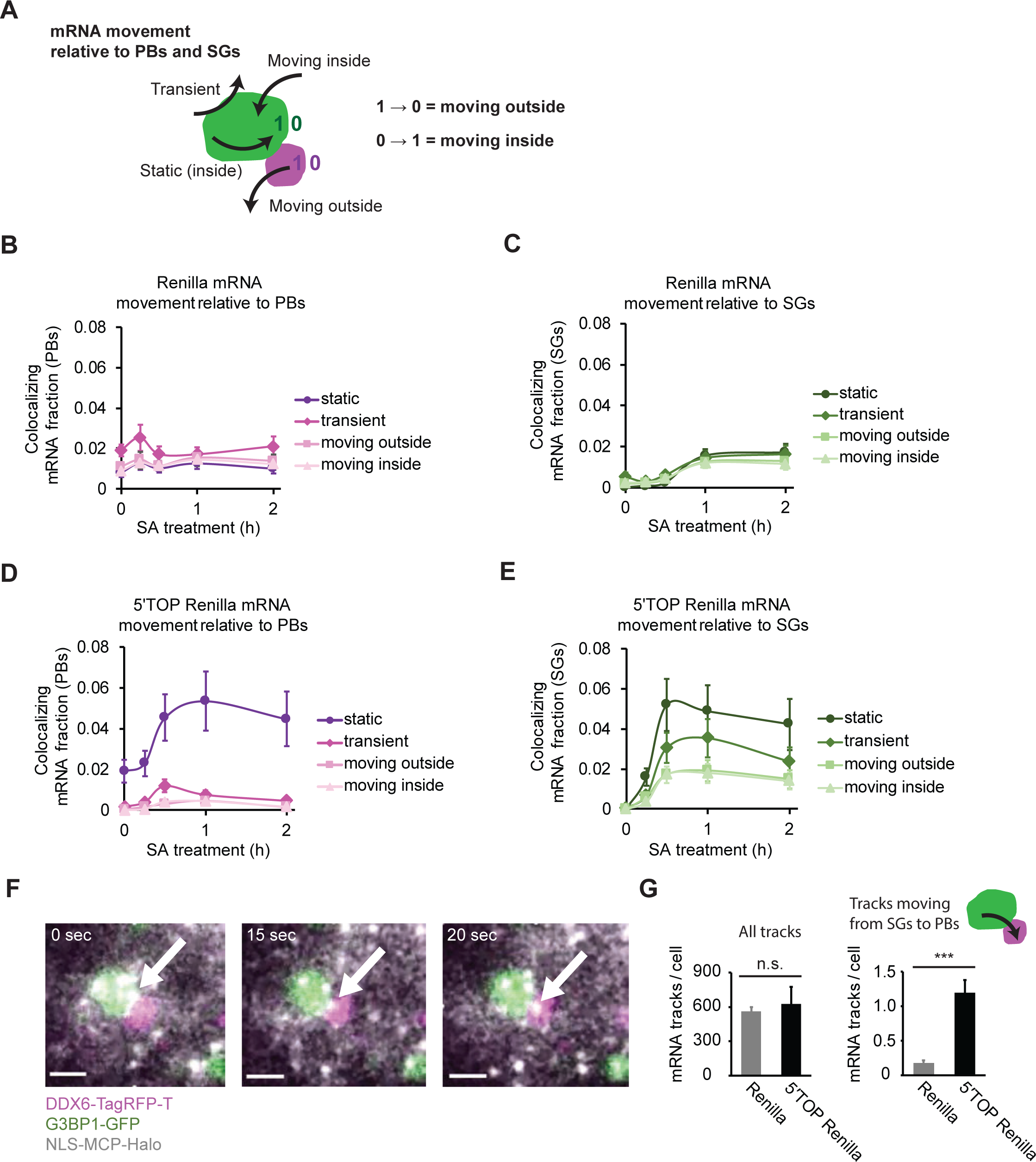
mRNA tracking reveals recruitment dynamics into SGs and PBs. (A) Data analysis workflow to quantify the movement of mRNAs relative to PBs and SGs. A localization index change from 1 to 0 represented an outward movement, a change from 0 to 1 represented an inward movement relative to a PB or SG. mRNA tracks with localization indices of exclusively 1, were considered to be static. Tracks with more than one entry and exit event were categorized as transient interactions. (B,C) HeLa cells stably expressing G3BP1-GFP, DDX6-TagRFP-T, NLS-MCP-Halo coat proteins and Renilla reporter mRNAs were treated with 100 µM SA for 2 hours and cells were imaged over time and their mRNA movement patterns were analyzed. Renilla mRNAs had no predominant movement pattern relative to PBs (B) or SGs (C) during the stress time-course. (D,E) HeLa cells stably expressing G3BP1-GFP, DDX6-TagRFP-T, NLS-MCP-Halo coat proteins and 5′TOP Renilla reporter mRNAs were treated with 100 µM SA for 2 hours and cells were imaged over time and their mRNA movement patterns were analyzed. (D) PB-associated mRNAs were mostly static. (E) SG-associated mRNAs were mostly static or showed transient interactions. (F) 5′TOP Renilla reporter mRNAs can move from a SG to a PB during SA stress. (G) Analysis of all mRNA movement patterns for shuttling events from SGs to PBs indicated that only a minor fraction of cytosolic 5′TOP Renilla reporter mRNAs move between both granules. Scale bars = 3 µm; mean ± SEM; two-tailed, unpaired Student′s t-test; *** = p < 0.001; > 20 fields of view per time point and experiment, 3 biological replicates.

The time course experiments indicated that mRNA recruitment to granules correlates with granule size and number and that there is a significant amount of mRNA exchange between granules and the cytosol during earlier and later phases of stress (fig. S1, Fig. 3B-E). Since SGs and PBs have been found to interact frequently and dynamically with each other (*31*), it has also been proposed that mRNAs can be sorted from SGs to PBs in a process referred to as “mRNA triage” (*32, 33*). We specifically searched for mRNA tracks within our stress time course data set that moved directly from SGs to PBs and were able to detect a small number of such events (Fig. 3F). The frequency of these events across the entire duration of the 120-minute time course was, however, extremely low. For, on average, ∼600 detected mRNA tracks per cell we could only identify 1 event using the 5′TOP Renilla reporter and 0.2 events for the Renilla reporter (Fig. 3G and Movie S7). We also searched for mRNA movement events in the inverse direction from PBs to SGs, but were not able to detect such events. Presumably, this is due to the high static mRNA localization and low outside mRNA movement rates of PBs (Fig. 3B).

To what extend the sequestration of mRNAs into granules has an effect on their decay and translation is currently unclear. Previously, we have found that translation and degradation of Renilla and 5′TOP Renilla mRNAs is globally inhibited throughout the cytosol, regardless of granule localization, during the stress response (*15, 34*). It has, however, been suggested that stress-induced PBs (*35, 36*) and SGs (*33*) could serve as sites for storage where mRNA molecules could be protected from the harmful effects of stress. Additionally, arsenite stress could result in the oxidation of mRNAs that can potentially lead to decreased mRNA half-lives through no-go decay (*37, 38*). While only ∼15% of 5′TOP Renilla mRNA reporters were found to be inside of PBs and SGs during stress, this provided an entry point for exploring the effect of this localization on the fate of transcripts during recovery from stress (Fig. 2).

To assess the potential protective effect of granule localization on mRNA decay, we first used3′-RNA End Accumulation during Turnover (TREAT) to quantify mRNA degradation with single-molecule resolution in unstressed cells (Fig. 4A) (*34*). In order to measure mRNA decay during recovery from stress, transcription was induced with doxycycline for 45 minutes followed by addition of SA for an additional 45 minutes and then cells were washed to remove both doxycycline and SA. Doxycycline removal stopped transcription of the mRNA reporter, so that only transcripts that experienced stress were monitored during the recovery phase. Cells were then fixed at different time points during an 8-hour stress recovery time course and intact and stabilized 3′-end fragments were quantified by smFISH (Fig. 4B). Interestingly, mRNA degradation remained inhibited for ∼2 hours after removal of SA despite the dephosphorylation of eIF2< and the dissolution of PBs and SGs during the first hour of recovery (Fig. 4B, fig. S1A, fig. S5). The single-molecule sensitivity of TREAT allows us to exclude the possibilities of either rapid mRNA decay of cytosolic or granule-localized transcripts during this initial period (fig. S6A). Using the TREAT data in stressed cells, we then tested several possible mRNA decay models that included a variable to account for the delay in mRNA degradation and allowed for the possibility for cytosolic and granule mRNAs to have different decay rates. The simplest model that reproduces our observed data had a delay of 2.2 ± 0.1 hours and a single decay rate for both cytosolic and granule mRNAs that was slightly faster than in unstressed cells (Fig. 4A, B and fig. S6B).

**Fig. 4:**
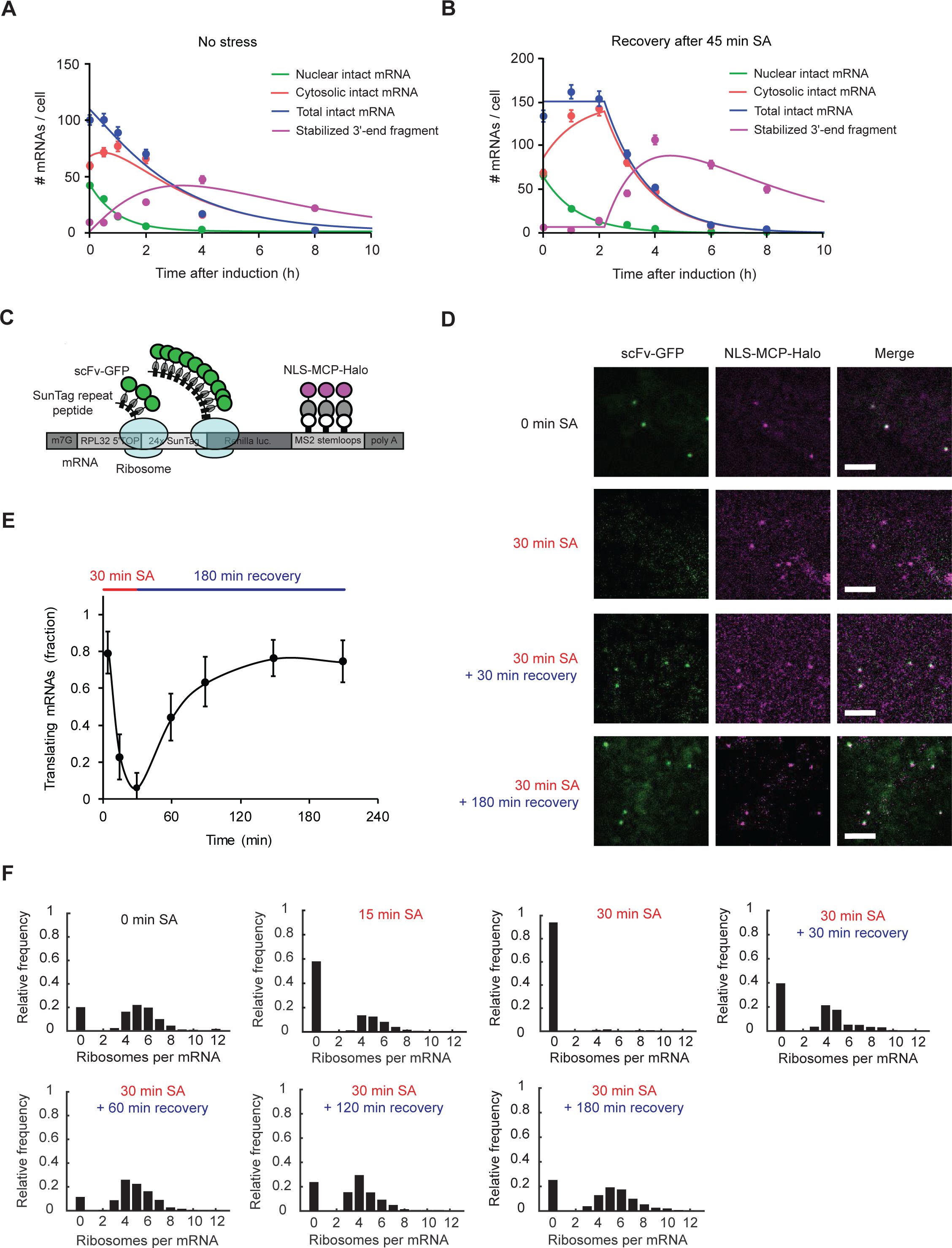
5′TOP mRNAs are not rapidly degraded and resume translation during recovery from stress. (A) TREAT measurement of mRNA decay in unstressed cells. Counts of intact mRNAs (nuclear and cytosolic) and stabilized 3′-ends were obtained from quantitative analysis of smFISH data at indicated time points (>200 cells per time point in two biological replicates). Data were fit to the model to calculate the nuclear export rate (1.2±0.2 h^-1^, R^2^=0.99), mRNA decay rate (0.4±0.1 h^-1^,R^2^=0.96) and stabilized 3′-end decay rate (0.4±0.2 h^-1^, R^2^=0.86). (B) TREAT measurement of mRNA decay in cells stressed with 100 µM SA. Counts of intact mRNAs (nuclear and cytosolic) and stabilized 3′-ends were obtained from quantitative analysis of smFISH data at indicated time points (>200 cells per time point in two biological replicates). Data were fit to the model to calculate the nuclear export rate (0.9±0.2 h^-1^, R^2^=0.99), mRNA decay rate (0.7±0.2 h^-1^,R^2^=0.98) and stabilized 3′-end decay rate (0.2±0.1 h^-1^, R=0.95) with a delay of 2.2±0.1h before decay resumes. (C) Schematic depiction of the 5′TOP SunTag Renilla mRNA reporter. Single-chain antibodies fused to GFP (scFv-GFP) label the ribosome emerging SunTag peptide chain in a length-dependent manner. (D) Representative images for SunTag translation imaging in cells stably expressing scFv-GFP, NLS-MCP-Halo, and inducible 5′TOP SunTag Renilla mRNA reporters. Under non-stress conditions most mRNAs (NLS-MCP-Halo) colocalized with a translation site (scFv-GFP). After 30 minutes of 100 µm SA treatment translation was blocked. During recovery from stress, translation sites colocalizing with mRNAs reappeared. (E) Quantification of the fraction of 5′TOP SunTag Renilla mRNAs colocalizing with translation sites showed that mRNA translation fully resumed to pre-stress levels during the recovery from stress. (F) The ribosomal occupancy distribution on mRNAs decreased during 30 minutes of SA treatment and reached a pre-stress distribution after 180 minutes of recovery from stress. Scale bar = 2 µm; mean ± SEM.

In order to determine the effect of granule localization on translation, we utilized a recently developed nascent polypeptide-based translation imaging system, since it offers the possibility to quantify the fraction of translating mRNAs per cell and their individual translational dynamics (*39–43*). This technique relies on the binding of single-chain antibodies fused to GFP (scFv-GFP) to the nascent SunTag epitopes that emerge from the ribosome and allows a fluorescence-based measurement of translation per mRNA molecule. We fused a 24x SunTag repeat cassette to the N-terminus of the Renilla luciferase coding sequence of our reporter, giving rise to a 5′TOP SunTag Renilla reporter (Fig. 4C). We then genomically integrated a single copy of this reporter into the previously used doxycycline-inducible HeLa cells. In addition, scFv-GFP was stably integrated into the cells. Individual mRNAs were visualized by the binding of NLS-MCP-Halo to the MS2 stem loops in the 3′UTR of the reporter.

We then used these cells to quantify the translation of individual mRNA molecules before, during and after stress. In the absence of stress, the majority of 5′TOP SunTag Renilla reporters (∼80%) were undergoing active translation as detected by the colocalization of the SunTag GFP signal with the NLS-MCP-Halo signal (Fig. 4D,E and Movie S8). After 30 minutes of SA-induced stress, almost all mRNAs (> 95%) were translationally inhibited, indicated by the absence of scFv-GFP labelled translation sites on mRNAs (Fig. 4D,E, and Movie S9). Next, we used the colocalization frequency of scFv-GFP with NLS-MCP-Halo to quantify the fraction of mRNAs undergoing active translation for all time points during the stress and recovery time course (Fig. 4E). If only the 15% of 5′TOP SunTag Renilla mRNAs bound to stress-induced mRNPs would be protected from stress, we expected that during translational recovery we should not observe more than 15% of mRNAs undergoing translation. Our experiment, however, indicates that 44% of all cytosolic mRNAs had already resumed translation after only 30 minutes of translational recovery and the fraction of mRNAs undergoing translation then gradually recovered over the next 2.5 hours to levels comparable to the pre-stress time point (Fig. 4E and Movie S10).

Due to the binary readout of using colocalization for the determination of translation, it remained a possibility that oxidative stress-inflicted chemical damage to non-sequestered mRNPs might decrease their translational efficiency. These potential defects in translation should be manifested in the number of ribosomes per mRNAs. SunTag-based translation imaging allowed quantifying the ribosomal occupancy per mRNA by dividing the fluorescent intensity of the translation site by the fluorescent intensity of a mature SunTag Renilla protein. We analyzed the distribution of all translation site intensities for all stress and recovery time points and calculated the ribosome occupancy per mRNA (Fig. 4F). In unstressed cells, each mRNA was bound by 4-5 ribosomes. During stress, the number of ribosomes per mRNA dropped after 15 minutes consistent with ribosomes running off until translation was almost completely inhibited at 30 minutes. After only 30 minutes of recovery when granules are still present in cells, the average ribosome occupancy increased to 3 ribosomes per mRNAs and after 3 hours of recovery, most mRNAs had regained their full ribosome occupancy and were bound by 4-5 ribosomes per mRNA (Fig. 4F). These results indicate that localization of an mRNA to granules during stress does not dramatically alter its translation when stress has been relieved.

We have characterized the dynamics of mRNA localization to SGs and PBs and its functional consequence during arsenite stress, however, granule composition and function may be altered when induced by other stresses or for disease-related granules (*7, 44–46*). The incorporation of LARP1 into granules and its anchoring of 5′TOP transcripts within them may provide an additional level of regulation of ribosome biogenesis in conditions when translation must be down-regulated (*47*). Since TIA-1 and TIAR are essential for SG formation and have also been shown to bind 5′TOP transcripts, future work will address how these interactions are coordinated during the stress response (*14*). Phase separated compartments are increasingly being observed in diverse biological contexts and are thought to organize intra-cellular biochemical reactions (*48*). Our work provides a framework for using single-molecule measurements to directly investigate molecular mechanisms within their cellular environment.

## Author Contributions

J.H.W. performed experiments and analyzed data with help from F.V. (image analysis) and I.H. (TREAT). Y. Z. and G.R. performed the mathematical modeling. J.H.W. and J. A. C. wrote the manuscript with input from all of the authors.

## Acknowledgements

This work was supported by the Novartis Research Foundation (J. A. C), the Swiss National Science Foundation grant 31003A_156477 (J.A.C), and the SNF-NCCR RNA & Disease (J.A.C). The authors thank K. Schönig (CIMH) for the parental HeLa 11ht cell line and L. Lavis (Janelia Farm) for providing Halo and SNAP dyes, T. Lionnet (Janelia Farm) for providing access to AIRLOCALIZE detection software, and J. Lykke-Andersen (UCSD) for sharing plasmids containing RPL32 5′TOP sequences. We acknowledge L. Gelman and S. Bourke (FMI) for microscopy support and H. Kohler (FMI) for cell sorting. We thank L. Giorgetti and all members of the Chao lab for their helpful discussions.

